# A platform for standardized, online delivered, clinically applicable neurocognitive assessment: WebNeuro

**DOI:** 10.1101/2023.08.28.553107

**Authors:** Leanne M. Williams

## Abstract

We are in the midst of a paradigm shift in which we see a precision mental health and the emergence of brain-based assessments for personalizing treatments. The precision emphasis of precision mental health includes a focus on measurement. Within the biopsychosocial domains of measurement, objective measures of cognitive function have an important role in disorders of mental health. Here, data integrating the development, norming, validation, psychometric testing and clinical application of a web-delivered neurocognitive assessment (WebNeuro) are presented. WebNeuro is designed for low-burden assessment of 12 domains of general and emotional cognition within 35-40 minutes, remotely and via each participant’s own computer. Data are from 1,317 healthy participants and over 500 clinical participants across diagnoses. Results provide norms across nine decades, and demonstrate the construct validity, internal consistency, and test-retest reliability of WebNeuro. Findings also show that WebNeuro identifies neurocognitive impairments across multiple neuropsychiatric disorders.

## INTRODUCTION

We are in the midst of a paradigm shift in which we see a precision mental health and the emergence of brain-based assessments for personalizing treatments. The ‘precision’ emphasis of precision mental health includes a focus on measurement. Within the biopsychosocial domains of measurement, objective measures of cognitive function have an important role in disorders of mental health. In the most prevalent and burdensome mental disorders, cognitive impairment is a major factor contributing to poor treatment outcomes, functional disability, suicide risk and lost productivity^1-15^. Researchers and practitioners in precision mental health require a scientifically sound and efficient to use platform for assessing cognitive function across general and emotional domains of function. Here, a cognitive assessment platform called Webneuro is described, with a synthesis of the robust norms across the lifespan, psychometric testing and application in clinical samples. This platform is delivered via a streamlined online system, enabling each individual to complete the assessment on their own local computer, within 35-40 minutes.

Development and testing of WebNeuro drew on advances in computer-driven behavioral testing platforms that utilizing the principles of neuropsychology within a digital format. The computerized format enables performance to be recorded in a consistently objective manner and for data to be quantified and normed in a standardized manner for each individual subject, and for studies, clinics and multi-center collaborations.

## METHODS

### Participants

A total of 1000 individuals (465 males, 535 females), aged 6-91 years (mean = 41.08, standard deviation = 23.04), were recruited in collaboration with the BRAINnet initiative. All participants gave written informed consent in accordance with local health and medical research council ethical guidelines. Data from 26 participants required exclusion due to an estimated IQ score of more than 1 standard deviation below the mean (i.e. less than 85). Therefore, the final sample consisted of 974 individuals (457 males, 517 females), aged 6-91 years (mean = 40.74, standard deviation = 23.10).

Web-based questionnaires ensured inclusion criteria were met: hearing or vision within normal (or corrected to normal) range, and estimated IQ (based on Spot the Word test;^16^) within normal range. Exclusion criteria were symptoms of an Axis 1 disorder determined using the BRID personal history and screening assessments (including SPHERE^22^) and Patient Health Questionnaire (PHQ9^17^) and items to screen for family history of psychiatric disorder (defined in terms of severity requiring medication and/or hospitalization). Web-based assessments also screened for physical brain injury (causing loss of consciousness for 10 minutes or more), neurological disorder, other serious medical or genetic condition and drug dependence, using the Alcohol Use Disorders Identification Test of the WHO (AUDIT) and Fagerstrom Tobacco Dependency Questionnaire^18^.

### Age distribution

Participants spanned ten decades of the human lifespan, with the following age and sex distributions: 6-9 years (*n* = 83, males (M): 38, females (F): 45), 10-19 years (*n* = 163, M: 90, F: 73), 20-29 years (*n* = 176, M: 65, F: 111), 30-39 years (*n* =76, M: 31, F: 45), 40-49 years (*n* = 56, M: 25, F: 31), 50-59 years (*n* = 60, M: 21, F: 39), 60-69 years (*n* = 276, M: 136, F: 140), 70-79 years (*n* = 74, M: 45, F: 29), 80-91 years (*n* = 10, M: 6, F: 4).

### Expanded norm cohort

To develop norms the original sample of 1000 was expanded further to create a normative cohort totaling 1,317 healthy participants (51% females), with an age range of 6 to 92 years (Mean=38 years, SD=20.6).

### Cognitive Assessments

WebNeuro is a fully computerized battery of tests designed for ready application in clinical research and clinical settings. It can be delivered remotely for each participant to complete on their own local computer. In addition, WebNeuro can be administered on a dedicated computer in a controlled setting as required by different protocols or clinical settings. WebNeuro has been validated (correlating.86 overall) against an original touchscreen version of the same tests^19^, and against traditional paper-and-pencil neuropsychological tests. WebNeuro contains tests of both emotion processing and general cognition^20,21^, as described below.

### Emotional Cognition

The Facial Expressions of Emotion Test (FEET) comprises two components: immediate explicit emotion identification and delayed implicit emotion priming^21^. Emotional expression stimuli comprise 16 different individuals (8 females, 8 males), each depicting neutral and evoked happiness, fear, sadness, anger, and disgust, adapted from the standardized set of Gur et al.^22^. Stimuli were adapted to be equated in terms of size, luminance, and central alignment of the face within the image (with eyes as midpoint reference).

#### FEET Explicit emotion identification

This condition requires explicit emotion identification via labeling of the expression. A total of 48 face stimuli (8 different individuals, depicting the 6 expressions) were presented in a pseudorandom sequence, for 2 seconds each, and participants identified (via computer mouse click) the verbal label for each emotional expression from among the six expression options. Accuracy, reaction time (RT) for correct responses and variability of RT for these responses were recorded. For incorrect responses, the expression selected on these trials was also catalogued.

#### FEET Implicit emotion priming

In this condition, the automatic implicit effects of emotion on general face recognition is assessed. Following an interval of 20 minutes, in which unrelated filler tasks were completed, the implicit emotion condition was presented. The original list of 48 expression stimuli is again presented in a pseudorandom sequence, but in this case each face is presented with a second ‘new’ face from the remaining 48 stimuli. These new faces were matched to each original face in terms of sex, approximate age, and expression. Participants identify which of the two faces they recognize from the original list presented under the emotion identification condition. Accuracy of face recognition is assessed to determine completion of the task. Reaction time for the implicit priming effect of each emotion (minus neutral) is calculated.

### General Cognition

#### Motor Tapping

Participants are required to tap the space bar with their index finger as fast as possible for 60 seconds, first with dominant and then non-dominant hand. Dependent measures are number of finger taps, as well as mean and variability (in standard deviations) of reaction time.

#### Choice Reaction Time

One of four target circles is luminated in a pseudorandom sequence, and participants are required to touch the luminated circle as quickly as possible following presentation. Twenty trials are administered with a random delay between trials of 2-4s.

#### Verbal Memory

A test list of 20 English words is presented over three trials, with immediate recognition examined at each trial. Memory recognition is assessed by presenting each word from the list alongside two distractor words from a new list of 40 words, and the order of presentation is pseudorandomized. This test assesses constructs equivalent to those assessed by the California Verbal Learning Test.

#### Digit Span

A series of digits (e.g., 4, 2, 7 etc., 500ms presentation) is presented and participants are asked to recall the order of digits on a numeric keypad; in both forward and reverse order trials. The number of digits in each sequence is gradually increased from 3 to 9, reflecting increasing difficulty. This test is equivalent to neuropsychological tests of digit span forwards and backwards.

#### Verbal Interference

Colored words with incongruent color-word combinations were presented and participants were first required to identify the name of each word as quickly as possible, and then to name the color of each word as quickly as possible. This task is a web-based variant assessing similar constructs to Golden’s Stroop test.

#### Switching of Attention

A pattern of 13 numbers (1-13) and 12 letters (A-L) were presented and participants were required to touch numbers and letters alternatively in ascending sequence (i.e., 1 A 2 B 3 C…), as a web–based adaption assessing similar constructs to Trails Making B.

#### Go-NoGo

Participants response *only* to Go stimuli (the word ‘press’ in green) and to immediately inhibit responses to NoGo stimuli (the word ‘press’ in red). Speed and accuracy are stressed equally. Stimulus duration is 200ms with ISI 2.5s. Mean reaction time, total errors, and errors of commission (responding to red stimuli) and omission (failing to respond to green) are measured.

#### Delayed Memory

Following an interval of 20 minutes filled with unrelated tasks, the original 20 words from the Verbal Memory Test are presented with 40 new distractor words. The original words are presented one at a time, with 2 distractor words, to assess delayed recognition memory.

#### N-Back test

An N-Back test (also known as the Continuous Performance Test) in which participants respond to a consecutive letter (via space bar) in a series of letters (B, C, D and G; 200ms presentation; ISI 2.5s in the task instructions. Mean reaction time, total errors, and errors of commission (identifying a non-consecutive letter as a target) and omission (failing to identify a target letter as a target) are measured.

#### Executive Maze

Participants are required to identify the hidden path (requiring 24 correct moves) through an 8×8 grid by using arrow keys. Incorrect moves are indicated by one tone (and red cross) and correct by a different tone and green tick. Task completion relies on two consecutive correct maze completions (or 7min time out). Dependent measures are completion time, overrun errors and total errors.

These tasks in total take approximately 30 to 45 minutes to complete. The testing platform controls the testing interface so that test takers cannot access other programs, including web searches.

## RESULTS

### Norms and norm development

Norms have been established across 9 decades for each of the tests^20,21^.

The distribution of demographic variables (age, years of education, height, weight), for each sex across age in the normative sample are shown in Figure 1.

**Figure 1.**
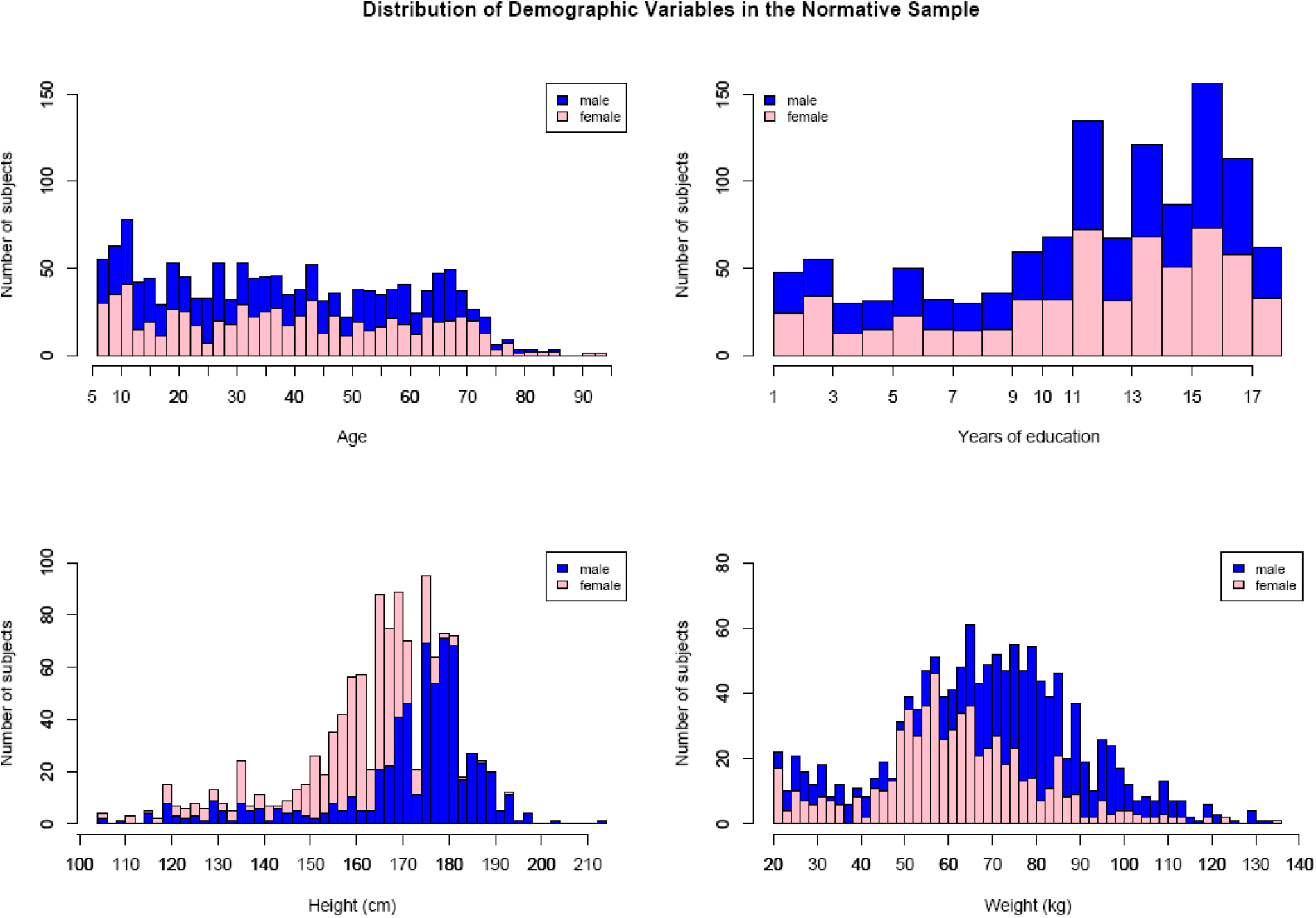
The distribution of demographic variables for the normative cohort (*n*=1,1317).

Norm data are plotted as Peer Regression Modeling (PRM) norm curves by sex and age band with 95% confidence intervals. Means and standard deviations (SDs) by sex and age band, separately for adults and minors, are then used as the reference data for new datasets.

The PRM technique is based upon and undertaken using the Crawford and Howell normalization procedure^23^. It allows the benchmarking of an individual’s scores so that subjects can be compared on individual differences over and above their differences in age, gender and education. PRM scores show how subjects relate to their healthy peers, which is especially relevant for distinguishing clinical groups from healthy controls. The PRM model used includes linear and logarithmic terms for ‘age’ and a linear term for ‘gender’. This expected score is subtracted from the subject’s actual score and the resulting difference is then divided by the standard error of the estimate of the regression equation. The result is a positive or negative number indicating the direction and magnitude of a subjects’ deviation from their age/gender predicted value. For example, a normalized score of 1.93 would reflect above average performance (in comparison to peers) and −2.23 would reflect below average performance.

## Reliability

### Test-retest reliability

148 participants (76 females, 72 males) ranging in age from 6 to 91 (Mean age=35.5 years, SD=16.9) were assessed twice with WebNeuro^24^. Analyses using the interclass coefficient method were used to establish the test-retest reliability of test performance.

Resulting reliability coefficients are shown in Table 1.

**Table 1.** Test-retest reliability of Self Regulation, Thinking and Emotion markers. Data are shown for subjects across all ages^1^.

| Domain and Test | Test-retest reliability |
| --- | --- |
| **Emotional Cognition** |  |
| Emotion Identification | 0.71 |
| Implicit Emotion Bias | 0.71 |
| **General Cognition** |  |
| Motor Tapping | 0.87 |
| Choice Reaction Time | 0.61 |
| Verbal Memory | 0.37 |
| Digit Span | 0.75 |
| Verbal Interference | 0.61 |
| Switching of Attention | 0.70 |
| Go-No-Go | 0.75 |
| N-Back | 0.71 |
| Maze | 0.68 |

### Multiple Forms

To avoid practice effects there are six parallel forms of the Webneuro tests. **Cross-cultural consistency** has also been established^25^.

### Face validity

The use of computerized assessments has a number of practical advantages. The American Psychological Association (APA) recognizes the value of computerized psychological testing and in 1987 published guidelines to assist in the development and interpretation of results from computerized test batteries (APA, 1987). Six major benefits of computerized assessment were identified:

1. Automation of data collection and storage
2. Greater efficiency of use
3. Freeing the clinician from test administration, enabling more focus on treatment
4. Increased sense of mastery and control for the client
5. Reduced negative self-evaluation among clients that experience difficulty on the computer
6. Greater ability to measure performance parameters that are not easily acquired through traditional paper-and-pencil tests (e.g., response latency, strength, variability).

WebNeuro is based on well-established constructs with face validity. Modifications to ensure face validity have been made in accordance with feedback from expert scientists and key opinion leaders. These developments have included attention to the ease of use, by a wide range of people. For example there are both adult and child wording versions of the instructions.

### Construct validity

PCA was used to evaluate the construct validity of the tests. The results are depicted as an associated pattern matrix illustrated in Tables 2 and 3.

**Table 2.**
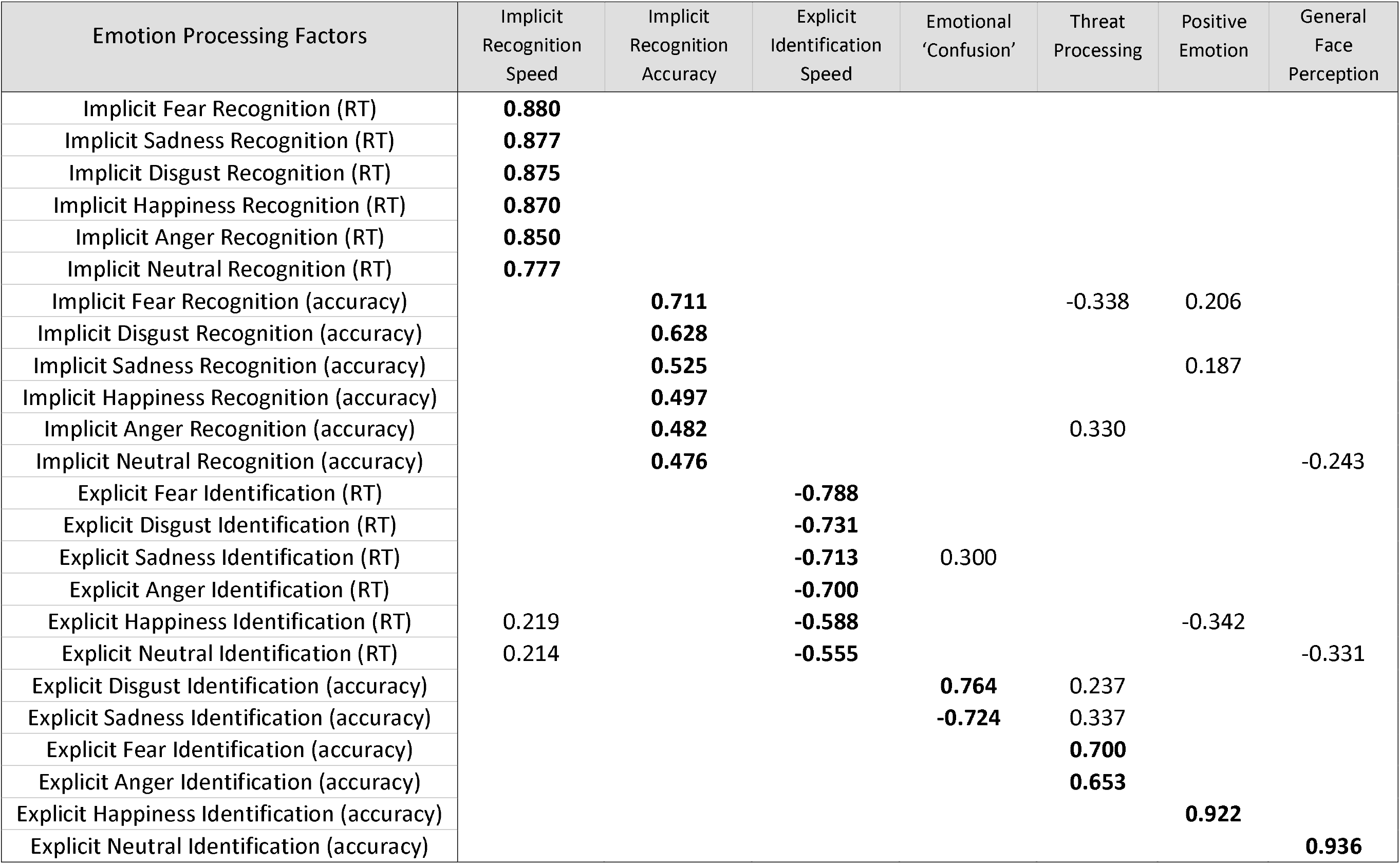
Factor loadings for the emotional cognition factors. Those highlighted in bold formed the seven factors, as labeled.

**Table 3.** Factor loadings for the general cognitive factors. Those highlighted in bold were used to form the seven factors, as labeled.

| General Cognitive Factors | Information<br>Processing<br>Speed | Inhibition/<br>Impulsivity | Working<br>Memory<br>Capacity | Verbal<br>Memory<br>Recognition | Sensori-Motor<br>Function | Executive<br>Function | Vigilance/<br>Sustained<br>Attention |
| --- | --- | --- | --- | --- | --- | --- | --- |
| Verbal Interference, name color (total score) | **-0.740** |  |  |  |  |  | 0.192 |
| Choice RT (average RT) | **0.658** |  |  |  |  |  |  |
| Verbal Interference, name word (total score) | **-0.605** |  |  |  |  |  |  |
| Switching of Attention (duration) | **0.552** |  | -0.199 |  |  |  |  |
| Go-NoGo (errors of commission) |  | **-0.886** |  |  | 0.181 |  |  |
| Go-NoGo (RT) |  | **0.849** |  |  |  |  |  |
| Digit Span Forward (max. number correct) |  |  | **1.002** |  |  |  |  |
| Digit Span Forward (total score) |  |  | **0.990** |  |  |  |  |
| Delayed Memory (total score) |  |  |  | **0.893** |  |  |  |
| Verbal Memory (total score) |  |  |  | **0.866** |  |  |  |
| Motor Tapping, dominant hand (pause) | -0.222 |  |  |  | **0.917** |  |  |
| Motor Tapping, dominant hand (total) | -0.278 |  |  |  | **-0.703** |  |  |
| Maze (overruns) |  |  |  |  |  | **0.936** |  |
| Maze (completion time) | 0.216 |  |  |  |  | **0.851** |  |
| N-Back (errors of omission) |  |  |  |  |  |  | **0.859** |
| N-Back (RT) | 0.219 |  |  |  | 0.177 |  | **0.609** |

#### Clinical association validity in clinical groups

WebNeuro clinical association validity has been established in clinical groups with Diagnosis in for clinical groups with confirmed with consensus from clinicians using the following criteria and scales. The groups and sample sizes are summarized below.

Adults

1. Major Depressive Disorder (Depression) (n=128): Structured Clinical Interview (DSM-IV) and Hamilton Rating Scale for Depression (HAM-D) (Hamilton, 1960).
2. Post-Traumatic Stress Disorder, PTSD (n=69): Structured Clinical Interview (DSM-IV) and Clinician Administered PTSD Scale (CAPS) (Blake et al., 1995).
3. Anxiety (Panic Disorder) (n=53): MINI International Neuropsychiatric Interview: Panic Disorder module (Sheehan et al., 2006) and the Composite International Diagnostic Interview (CIDI-Auto 2.1).
4. First Episode Schizophrenia (FES) (n=54): Structured Clinical Interview (DSM-IV) and Positive and Negative Symptoms Scale (PANSS) (Kay et al., 1990).
5. Alzheimer’s Disease (AD) (n=25): Clinical Dementia Rating (CDR) (Morris, 1993) and the Mini-Mental State Examination (MMSE) (Cockrell & Folstein, 1988).
6. Mild Cognitive Impairment (MCI) (n=76): Mini-Mental State Examination (MMSE) (Cockrell & Folstein, 1988) and the criteria outlined by Peterson et al. (1999). Children/Adolescents
7. Attention Deficit Hyperactivity Disorder (ADHD) (n=350): Structured Clinical Interview (DSM-IV) and Conners’ Rating Scale (Conners et al., 1998).
8. Anorexia Nervosa (n=42): Structured Clinical Interview (DSM-IV). Other Conditions
9. Sleep Disorder (Obstructive Sleep Apnea) (n=50): Multivariate Apnea Prediction Index (MAPI) of greater than 0.50.

## DISCUSSION

The present findings provide an overview and psychometric foundation for the use of a standardized platform for computerized assessment of general and emotional cognitive functions, suited for use in clinical samples relative to reference data.

These findings establish a reliable and valid assessment for assessing general and emotional cognition, with lifespan norms, across a wide range of mental health conditions. The assessment maximizes coverage of multiple domains of cognition while also minimizing patient burden. It comprises test of 12 general and emotional cognitive domains, and the total assessment takes only 35-40 minutes. WebNeuro is administered via a streamlined computerized system, such that participants and patients can complete the assessment on their own computer. The system is designed to provide a structured assessment without interference from other operations that may be occuring on each individual’s computer.

Performance on each of the WebNeuro tests is generated in both raw performance scores and in norm-referenced scores. Performance measures enable assessment of group trends as well as individual differences within healthy and clinical populations. Norm referencing facilitates the identification of general and emotional cognitive impairments at the individual patient level, relative to matched healthy peers. Assessment of these impairments can also help stratify subgroups within and across multiple diagnoses, and aid in the selection and management of tailored treatments.

## Footnotes

1 *Note: Test-retest reliability is truncated for this measure as there is a strong ceiling effect in healthy controls causing a restriction of range in **inter**-subject variability which artificially magnifies **intra**-subject variability in the correlation. The ceiling effect for controls is a trade-off that increases the sensitivity of the measure for detecting degree of memory deficits.

## REFERENCES

1. Ahmed, A.O., et al. Do cognitive deficits predict negative emotionality and aggression in schizophrenia? Psychiatry Res 259, 350–357 (2018).

2. Cole, M.W., Repovš, G. & Anticevic, A. The frontoparietal control system: a central role in mental health. Neuroscientist 20, 652–664 (2014).

3. Etkin, A., Gyurak, A. & O’Hara, R. A neurobiological approach to the cognitive deficits of psychiatric disorders. Dialogues Clin Neurosci 15, 419–429 (2013).

4. Fava, M., et al. A cross-sectional study of the prevalence of cognitive and physical symptoms during long-term antidepressant treatment. J Clin Psychiatry 67, 1754–1759 (2006).

5. Fleming, S.K., Blasey, C. & Schatzberg, A.F. Neuropsychological correlates of psychotic features in major depressive disorders: a review and meta-analysis. J Psychiatr Res 38, 27–35 (2004).

6. Gupta, M., et al. Relationships among neurocognition, symptoms, and functioning in treatment-resistant depression. Arch Clin Neuropsychol 28, 272–281 (2013).

7. Hack, L., Tozzi, L, Zenteno, S, Olmsted, A, Hilton, R, Yesavage J, Williams LM. A Cognitive Biotype of Depression Linking Symptoms, Behavior Measures, Neural Circuits, and Treatment Outcomes. Biological Psychiatry 93, S72–S73 (2023).

8. Hasselbalch, B.J., Knorr, U. & Kessing, L.V. Cognitive impairment in the remitted state of unipolar depressive disorder: a systematic review. J Affect Disord 134, 20–31 (2011).

9. Lam, R.W., Kennedy, S.H., Mclntyre, R.S. & Khullar, A. Cognitive dysfunction in major depressive disorder: effects on psychosocial functioning and implications for treatment. Can J Psychiatry 59, 649–654 (2014).

10. McIntyre, R.S., et al. The impact of cognitive impairment on perceived workforce performance: results from the International Mood Disorders Collaborative Project. Compr Psychiatry 56, 279–282 (2015).

11. Pu, S., Setoyama, S. & Noda, T. Association between cognitive deficits and suicidal ideation in patients with major depressive disorder. Sci Rep 7, 11637 (2017).

12. Rogers, M.A., et al. Executive and prefrontal dysfunction in unipolar depression: a review of neuropsychological and imaging evidence. Neurosci Res 50, 1–11 (2004).

13. Schatzberg, A.F., et al. Neuropsychological deficits in psychotic versus nonpsychotic major depression and no mental illness. Am J Psychiatry 157, 1095–1100 (2000).

14. Xiang, X. & An, R. The Impact of Cognitive Impairment and Comorbid Depression on Disability, Health Care Utilization, and Costs. Psychiatr Serv 66, 1245–1248 (2015).

15. Schussler-Fiorenza Rose, S.M., et al. Depression, health comorbidities, cognitive symptoms and their functional impact: Not just a geriatric problem. J Psychiatr Res 139, 185–192 (2021).

16. Baddeley, A., Emslie, H. & Nimmo-Smith, I. The Spot-the-Word test: a robust estimate of verbal intelligence based on lexical decision. Br J Clin Psychol 32, 55–65 (1993).

17. Kroenke, K., Spitzer, R.L. & Williams, J.B. The PHQ-9: validity of a brief depression severity measure. J Gen Intern Med 16, 606–613 (2001).

18. Heatherton, T.F., Kozlowski, L.T., Frecker, R.C. & Fagerstrom, K.O. The Fagerstrom Test for Nicotine Dependence: a revision of the Fagerstrom Tolerance Questionnaire. Br J Addict 86, 1119–1127 (1991).

19. Silverstein, S.M., et al. Development and validation of a World-Wide-Web-based neurocognitive assessment battery: WebNeuro. Behav Res Methods 39, 940–949 (2007).

20. Mathersul, D., et al. Explicit identification and implicit recognition of facial emotions: II. Core domains and relationships with general cognition. J Clin Exp Neuropsychol 31, 278–291 (2009).

21. Williams, L.M., et al. Explicit identification and implicit recognition of facial emotions: I. Age effects in males and females across 10 decades. J Clin Exp Neuropsychol 31, 257–277 (2009).

22. Gur, R.C., et al. A method for obtaining 3-dimensional facial expressions and its standardization for use in neurocognitive studies. J Neurosci Methods 115, 137–143 (2002).

23. Crawford, J.R. & Howell, D.C. Regression equations in clinical neuropsychology: an evaluation of statistical methods for comparing predicted and obtained scores. J Clin Exp Neuropsychol 20, 755–762 (1998).

24. Williams, L.M., et al. The test-retest reliability of a standardized neurocognitive and neurophysiological test battery: “neuromarker”. Int J Neurosci 115, 1605–1630 (2005).

25. Paul, R.H., et al. Cross-cultural assessment of neuropsychological performance and electrical brain function measures: additional validation of an international brain database. Int J Neurosci 117, 549–568 (2007).

